# Automatic segmentation and cardiac mechanics analysis of evolving zebrafish using deep-learning

**DOI:** 10.1101/2021.02.21.432186

**Authors:** Bohan Zhang, Kristofor Pas, Toluwani Ijaseun, Hung Cao, Peng Fei, Juhyun Lee

**Affiliations:** Joint Department of Bioengineering, UT Arlington/ UT Southwestern, Arlington, TX, USA, 76010; School of Optical and Electronic Information-Wuhan National Laboratory for Optoelectronics, Huazhong University of Science and Technology, Wuhan, China, 430074; Department of Electrical Engineering and Computer Science, UC Irvine, Irvine, CA, USA, 92697

**Keywords:** U-net, LSFM, segmentation, zebrafish, cardiac mechanics

## Abstract

**Objective:** In the study of early cardiac development, it is important to acquire accurate volume changes of the heart chambers. Although advanced imaging techniques, such as light-sheet fluorescent microscopy (LSFM), provide an accurate procedure for analyzing the structure of the heart, rapid and robust segmentation is required to reduce laborious time and accurately quantify developmental cardiac mechanics.

**Methods:** The traditional biomedical analysis involving segmentation of the intracardiac volume is usually carried out manually, presenting bottlenecks due to enormous data volume at high axial resolution. Our advanced deep-learning techniques provide a robust method to segment the volume within a few minutes. Our U-net based segmentation adopted manually segmented intracardiac volume changes as training data and produced the other LSFM zebrafish cardiac motion images automatically.

**Results:** Three cardiac cycles from 2 days post fertilization (dpf) to 5 dpf were successfully segmented by our U-net based network providing volume changes over time. In addition to understanding the cardiac function for each of the two chambers, the ventricle and atrium were separated by 3D erode morphology methods. Therefore, cardiac mechanical properties were measured rapidly and demonstrated incremental volume changes of both chambers separately. Interestingly, stroke volume (SV) remains similar in the atrium while that of the ventricle increases SV gradually.

**Conclusion:** Our U-net based segmentation provides a delicate method to segment the intricate inner volume of zebrafish heart during development; thus providing an accurate, robust and efficient algorithm to accelerate cardiac research by bypassing the labor-intensive task as well as improving the consistency in the results.

## 1. Introduction

Biomechanical analysis is important during cardiac development as assessment of biomechanics is closely associated with regulation of valve formation, ventricular septum, and trabecular morphology related to cardiogenic transcriptional and growth/differentiation factors [1; 2]. Lack of intracardiac biomechanical force could induce the malfunction of genetic programming resulting in congenital heart defects in humans and mice [3]. Volume change-based cardiac mechanics measurements (*e.g.* ejection fraction), from the complex trabeculated and beating heart, are most commonly used, and play an important role to evaluate the cardiac health condition. Such measurement relies on the accurate reconstruction of the heart’s volume, which in turn depends on the accurate segmentation of the biomedical images. Although the segmentation of biomedical images for volume reconstruction has been extensively studied in the radiation imaging techniques such as MRI, or CT[4; 5; 6], these imaging methods are challenging to be adopted for optical fluorescent images. Although various optical microscopes have been extensively used to study in biomedical research due to inexpensive approach, high resolution, and amenable fluorescent tagging [7; 8; 9], it suffers from light scattering and different intensity of fluorescent signal. In addition, imaging dynamic samples is another challenge for conventional microscopes. Unlike microscopes, however, light-sheet fluorescent microscopy (LSFM) circumvents these challenges to enable to capture *in vivo* dynamic sample, such as zebrafish heart, with high axial resolution, deep axial scanning, fast image acquisition, and low photobleaching [10; 11].

Despite having a two-chambered heart and a lack of a pulmonary system, the zebrafish represents an emerging vertebrate model for studying developmental biology [12; 13]. Its transparency and short organ developmental timeline enable rapid and high throughput analysis of developmental stages with optical fluorescent technology [14]. Such advantages of zebrafish and LSFM system make a powerful tool for the studying of *in vivo* cardiac development.

To understand the cardiac function and mechanics as well as further analysis, intracardiac segmentation is a necessary step [2; 15]. Previously, segmentation of the LSFM images for measuring cardiac mechanics was accomplished manually by recognizing different intensities from large amounts of samples or tissue scatterings engenders many variables [16]. The time consuming task of manually segmenting the LSFM images is infeasible when processing high axial resolution data, as the number of images required is enormous. On the other hand, lower axial resolution degrades the accuracy of the volume measurement, while inconsistent manual segmentation poses a threat to the overall quality of the cardiac mechanics analysis. Recently, Akerberg *et. al,* nicely demonstrated SegNet based deep-learning segmentation of zebrafish hearts at the early ventricular developmental stage before trabeculation [17]. However, segmentation of complex ventricular morphology after initiating trabeculation hampers accurate cardiac mechanics analysis. Therefore, we utilize the advancements in a specific convolution neural network (CNN) architecture, namely U-net [18], which performs binary classification of the pixels in the LSFM images. Unlike natural images, in which rich color and texture information are provided, LSFM images offer limited information. To reliably segment an LSFM image, a pixel location has to be considered. Our proposed U-net utilizes such information via a multi-scale processing pipeline, which uses the global branch to help locate a pixel while using the local branch to examine the neighborhood of the pixel. The main contribution of this paper is to propose and demonstrate a practical and robust U-net architecture for optical imaging, which is tailored to segment LSFM images of the developing zebrafish heart. Besides, the synchronization of the LSFM image sequence is applied before segmentation as a pre-processing step.

For our application, the U-net was trained to utilize LSFM images of a zebrafish during ventricular development from 2 days post fertilization (dpf) to 5 dpf. In this paper, we explore the potential use of the U-Net architecture to expedite the segmentation of the intracardiac zebrafish heart, including atrium and ventricle, and further biomechanical analysis of the extracted results from the network.

## 2. Methods

### Zebrafish preparation for imaging

The zebrafish used for this study was raised and maintained in our zebrafish core facility under required protocol under UT Arlington Institutional Animal Care and Use Committee (IACUC) protocol (#A17.014). Transgenic *tg(cmlc2:gfp)* zebrafish lines were used in this study to observe myocardium and chamber development. To ensure a clear image, a medium composed of 0.0025% phenylthiourea (PTU) was used to suppress pigmentation at 20 hours post-fertilization (hpf) [19]. Before imaging, zebrafish embryos were anesthetized in 0.05% Tricaine and immersed in a solution of 0.5% low melt-agarose at 37°C. The embryos were then transferred to a fluorinated ethylene propylene (FEP) tube (Refractive Index: 1.33) to minimize refraction along the path of light before image acquisition. This FEP tube was then immersed in water (Refractive Index: 1.33) and connected to an in-house LSFM to scan 500 images per slice from anterior to posterior of zebrafish heart with 2 μm thickness [20].

### 4D reconstruction of beating zebrafish heart

4D reconstruction of *in vivo* beating zebrafish heart was performed using previously described methods [16; 21]. Performed 4D reconstruction procedure is based on assumption that zebrafish heartbeat is regular throughout the image acquisition. However, experiments demonstrate that this assumption is not reliable. To minimize the natural irregularity of zebrafish heartbeat, we added an extra parameter (δ) to detect phase lock at period determination[16; 21] to observe irregularity during the experiments. This latent variable window, δ, can be set roughly −0.3 ms to +0.3 ms after finding the estimated cardiac period manually [22].

### Segmentation of intracardiac domain

Well reconstructed 4D images were selected for manual segmentation of the intracardiac domain. 3D images of each time point from the beating heart were loaded into Amira software (Thermofisher Scientific, Waltham, MA) or 3DSlicer. First, the inner cavity of each 2D slice was carefully selected manually and reconstruct 3D volume segmentation for each time point of the beating heart. We repeated this process at all developmental points to make segmentation masks. Manual segmentation masks were used for training our U-net architecture.

### Application of U-net architecture

The U-net architecture revolves around two distinct paths: the contracting path and the expansive path [18]. The network takes the input of size: A×A×n{A= 2k|k∈N}array and pushes it initially through the contracting path, which processes the input through a series of 3×3 convolutions, the application of a Rectified Linear Unit (ReLU) following each convolution, then a 2×2 max-pooling operation. This yields an output of A/2×A/2×2n, expanding the feature channel by double. This process is repeated four times before the final output begins being processed by the expansive path, which undergoes a similar process, with an exception of instead of max-pooling, the operation is a 2×2 up convolution (**Figure 1**).

**Figure 1.**
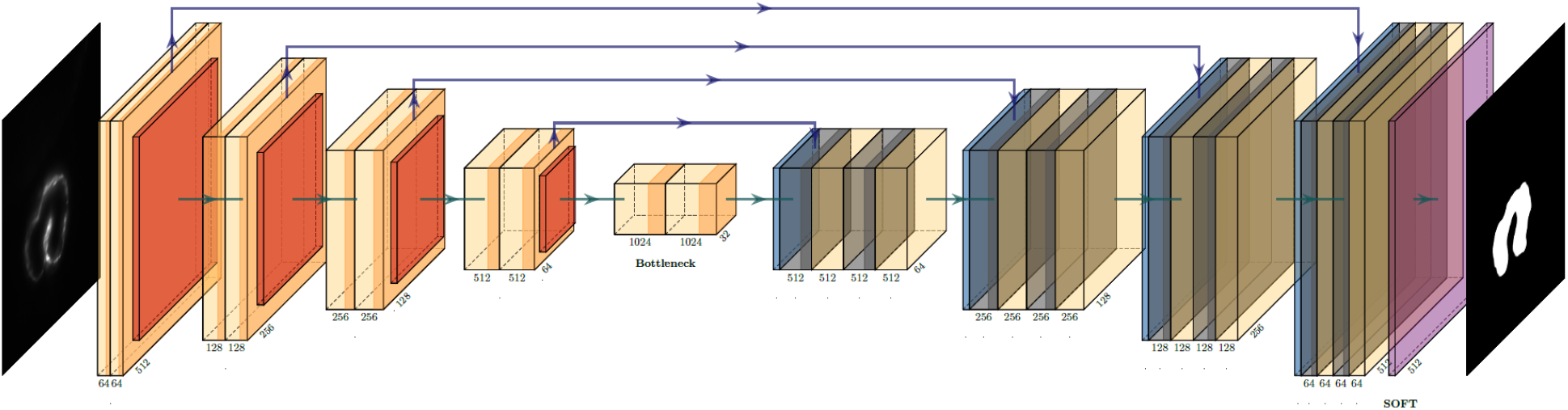
U-net CNN architecture utilized to generate the binary mask of the intracardiac domain of zebrafish. Each box represents a multi-channel feature map which allows for efficient and accurate extraction of anatomical features. In our specific application, the input was a 512×512 pixel map.

### Network training with manually segmented images

Training data in regard to CNN learning for this particular application is divided into two separate bins: training volumes and labels. The training volumes were acquired through the python program. For these, we selected entire slices of the 20 samples of 3D volume of heart, representing three cardiac cycles, with each slice containing n images (n∈N) which was dependent on the slice selected. Following data acquisition, hand-segmentation was performed using 3DSlicer GUI application on a 2D plane spanning the entirety of the zebrafish heart captured. Both these training bins were converted into 8-bit format and imported into U-NET as a 512×512×n8-bit array, with bijective correspondence between volumes and labels.

### Dice Similarity Coefficient Correlation

The primary method to which our automatic segmentation was assessed was using the Dice Similarity Coefficient Correlation, which compares the amount of space of which the volumes of the automatic and hand segmentation overlap in comparison with the summation of the total number of pixels. The calculation of this is as follows for area A, representing auto-segmentation and area B, representing our manually-labeled segmentation:

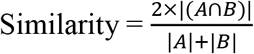

The outcome yields some value *x* for *x* ∈ [0,1] with the value one (approximating a near-perfect segmentation match with respect to the initial hand segmentation used) and zero (showing essentially no correlation between the two).

### Cardiac mechanics analysis

After segmented volume, we multiplied voxel resolution to obtain the actual volume of zebrafish heart. We capture the size of most dilated points and most contracted points as end-diastolic volume (EDV) and end-systolic volume (ESV), respectively. Stroke volume (SV) was calculated by subtracting ESV from EDV. Ejection fraction (EF) is the ratio of blood ejection which can be simply calculated from EF = (SV/EDV)×100.

## 3. Results

### Manual segmentation from LSFM for U-net training

Manual segmentation was performed along the 2D axial plane of the zebrafish heart for each studied sample. The methodology for manual segmentation, which served as our ground truth and labels, was performed using a contour-based segmentation method provided by 3DSlicer and was done in conjunction with ad hoc edits done on the resulting figures. Manually segmented images were used as training datasets for U-net (**Figure 1**). Concept of U-net architecture was to supplement a usual contracting network by successive layers, where pooling operators were replaced by upsampling operators [18]. Here, original 512×512 pixels LSFM images were deconstructed to detect features of intracardiac boundaries. Then upsampling of each layer increased the resolution of the output. To localize, higher resolution features from the downsampling path were combined with the upsampled output. As inputting manually segmented images to train, a successive convolution layer learned to assemble a more precise output based on this information. In addition, a Gaussian 2D filter of kernel size of 2.0 μm was applied to smooth manual segmentation. Contour interpolation was further applied to the 3D structure output to minimize the step size of later 3D reconstruction. The fidelity of the manual segmentation was done observationally through inspecting the spatial comparison between the label generated and the intracardiac area of the 2D slice (**Figure 2**).

**Figure 2.**
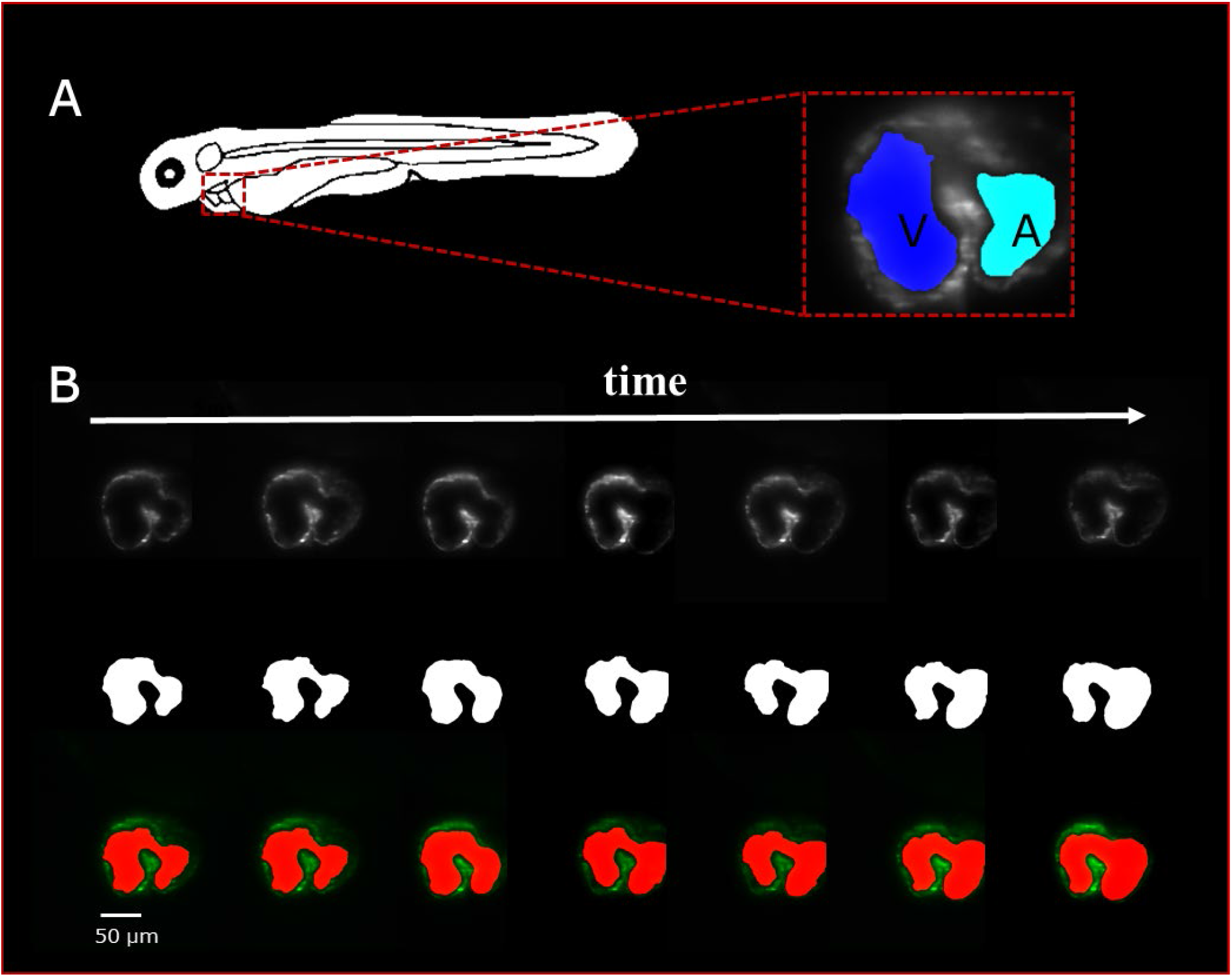
Sequence of selected LSFM images with the manual hand segmentation mask from 4 dpf zebrafish heart. (A) Diagram showing the anatomical feature of the zebrafish heart as well as a single axial slice with corresponding binary mask generated by the U-net. It is observed that there is a clear distinction between the atria and ventricle of the specimen. (B) A sequence of selected axial slices with corresponding binary masks generated by our U-net program, the reference scale bar is of size 50 μm.

### Validation of U-net based auto-segmented image

We visualize the segmentation ability of our networks by reconstructing the diastolic stage and systolic stage of zebrafish (**Figure 3A-B**). Although there is small segmentation discrepancy between ground truth and auto-segmentation around fluorescent boundaries (**Figure 3C-D**), U-net based-segmentation accuracy for each frame has a mean Dice coefficient score of 0.95 with standard deviation 0.02 (**Figure 3E**). Due to trabeculation in ventricle at 4 dpf, manual segmentation was more sophisticatedly segmented in rough fluorescent boundaries. Although our auto-segmentation performed to detect rough boundaries in trabeculated area, the roughness was relatively smooth. However, smooth surface in atrium and innermost area of ventricle captured more closely to the fluorescent signal.. This discrepancy could be from the dataset, that we trained the network, inconsistency of manual segmentation judged by human. Albeit, this demonstrates remarkable ability of our U-net based segmentation to segment intracardiac chamber with high dice coefficient score and reduces inherent manual segmentation.

**Figure 3.**
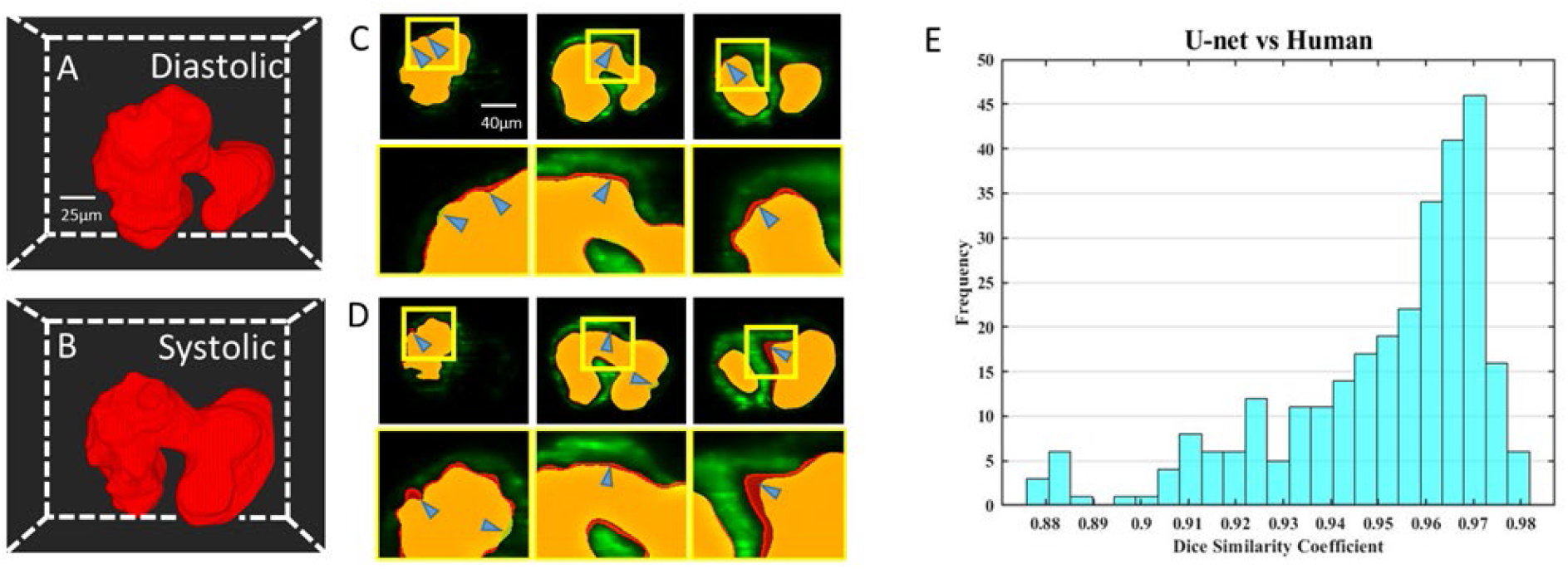
Comparison of U-net based auto-segmentation to manual hand segmentation. (A-B) Geometric representation of both the diastolic and systolic stages of the heart cycle. (C-D) Comparison in the 2D axial plane between auto-segmentation of the U-net and ground truth in both the diastolic and systolic stages respectively. The blue arrows indicate areas of disjoint space within the segmentations, shown in red (manual segmentation), green (auto-segmentation) and yellow (intersection). (E) Corresponding Dice Similarity Coefficient comparing the results between our ground truth and automatic segmentation demonstrated the ability of auto-segmentation.

### Automatic Morphology to Separate Inner Chamber Space

For the validation of the U-net-based auto-segmentation, to allow for further application, it was seen necessary to derive some function to separate the inner-chamber structure to allow for independent analysis of the two-volume components and their disjoint biomechanical characteristics. This was done using an in-house Matlab code utilizing an iterative process of applying an erosion operator relying on a 3D structuring element until fracturing the single master volume. Following this, a dilation operator was applied to the individual subvolumes to recover lost space, and the resulting dilated volumes were intersected in 3D space with the initial master volume. The output of this program was the resulting intersected space.

This program’s use proved to be critical in studying the resulting biomechanical elements of the atria and ventricle, as it allowed for consistency in determining the chambers (**Figure 4A).** The methodology was used successfully to distinguish the atrium and ventricle within the program on a three-dimensional level (**Figure 4B).** The resulting axial scans of the separated atria and ventricle show a clear distinction between the two different chamber areas per slice at 4 dpf (**Figure 4C).**

**Figure 4.**
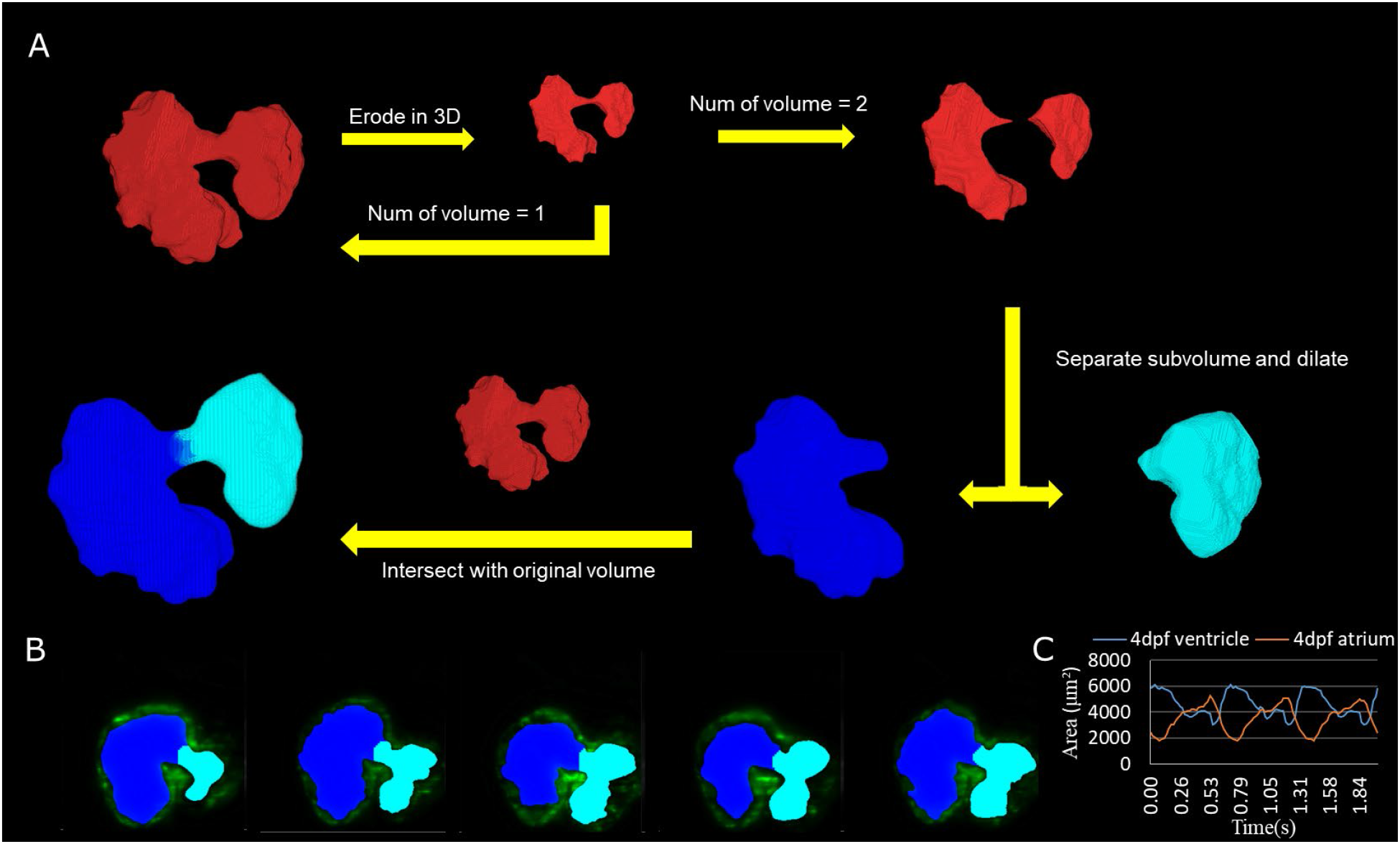
Subdivision of atrium and ventricle. (A) Methodology of using the automatic segmentation to define the morphology of inner-volume of the zebrafish heart. This method relies on an iterative process which uses an erosion operator on the 3D structure of the inner volume until division occurs. Following the division of the volume into two separate components, extraction of each individual piece follows with a dilation operation and intersection with the original volume. (B) 2D axial slices showing the differentiation of the Atrium (Teal) and Ventricle (Blue) of various slices. (C) Area change over time from 2D sliced (B) was successfully demonstrated.

### Volumetric analysis

We have applied 4D synchronization methods to reconstruct volume change over time [16; 20]. Analysis of the volume change along three cardiac cycles was provided to study the biomechanical changes over time (**Figure 5A**). Within this, the volume was plotted of the atria and ventricle independently and the total intracardiac volume of both chambers (**Figure 5B**). Although total cardiac volume of 4 dpf is oscillating during pumping, the mean value is roughly 9.5 × 10^5^ μm^3^. At 4 dpf, trabeculae developed in the ventricle; thus 4D reconstruction showed the motion of the corrugated surface of the ventricle. Due to lack of trabeculae [23], the atrium has relatively tranquil motion of curvature during atrial contraction and relaxation. Our results reveal the potential power of complex morphology and functional analysis.

**Figure 5.**
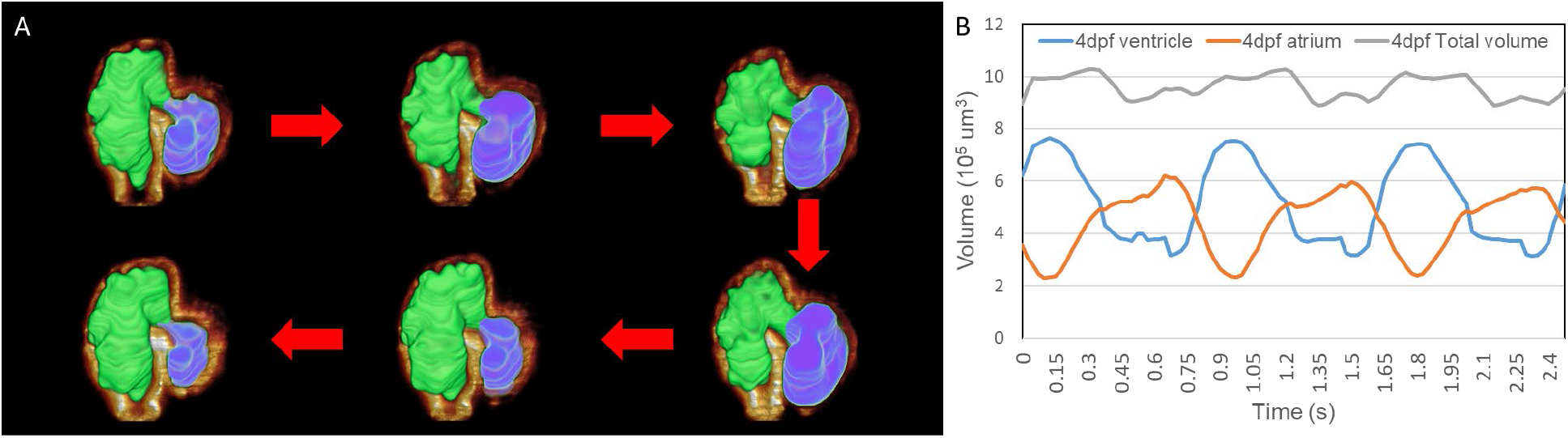
Representation of the contraction to dilation of zebrafish heart volume change over time. (A) Auto-segmented successfully reconstructed 4D image and captured the rough inner surface of zebrafish due to trabeculation after merging with fluorescently labelled *tg(cmlc2:gfp)* zebrafish from LSFM images. (B) Volume change of ventricle and atrium was measured from auto-segmentation. Total volume of atrium and ventricle represented around 9.5 × 10^5^ μm^3^ when zebrafish was at 4 dpf.

### Assessment of cardiac mechanics of developing zebrafish heart

In studying the volumetric change of the zebrafish heart during early-stage development between 2 dpf and 5 dpf, observations allowed for analysis of the atria and ventricle volumetric loads. Interestingly, we recognize that ventricular contraction pattern shifts left from 2 - 3 dpf while shifting right from 4 - 5 dpf. Similarly, a pattern was observed within the volumetric change of the atrium, albeit in an inverse manner. (**Figure 6A-D**). Furthermore, based on volume change data, we have performed cardiac mechanics analysis during development. We first analyze the end-diastolic volume (EDV) and end-systolic volume (ESV) of the atrium and ventricle. Although the trend of EDV and ESV of both atrium and ventricle consistently increased, there was a significant increase between 2 dpf to 3 dpf where morphology changed by cardiac looping [24] (**Figure 6E-F**). Also, the active trabeculation process[16; 23] which increases contractility of ventricle affects ESV of ventricle at 5 dpf from 4 dpf. Interestingly, the stroke volume (SV) of the ventricle consistently increased while the SV of atrium remains consistent around 3.6 × 10^5^ μm^3^ during early cardiogenesis (**Figure 6G**). We have further analyzed ejection fraction (EF) from EDV and ESV of the atrium and ventricle to understand how much blood each chamber pumps out with each contraction. Although, at 2 dpf, EF was high in both atrium and ventricle, it continuously decreased throughout development (**Figure 6H**).

**Figure 6.**
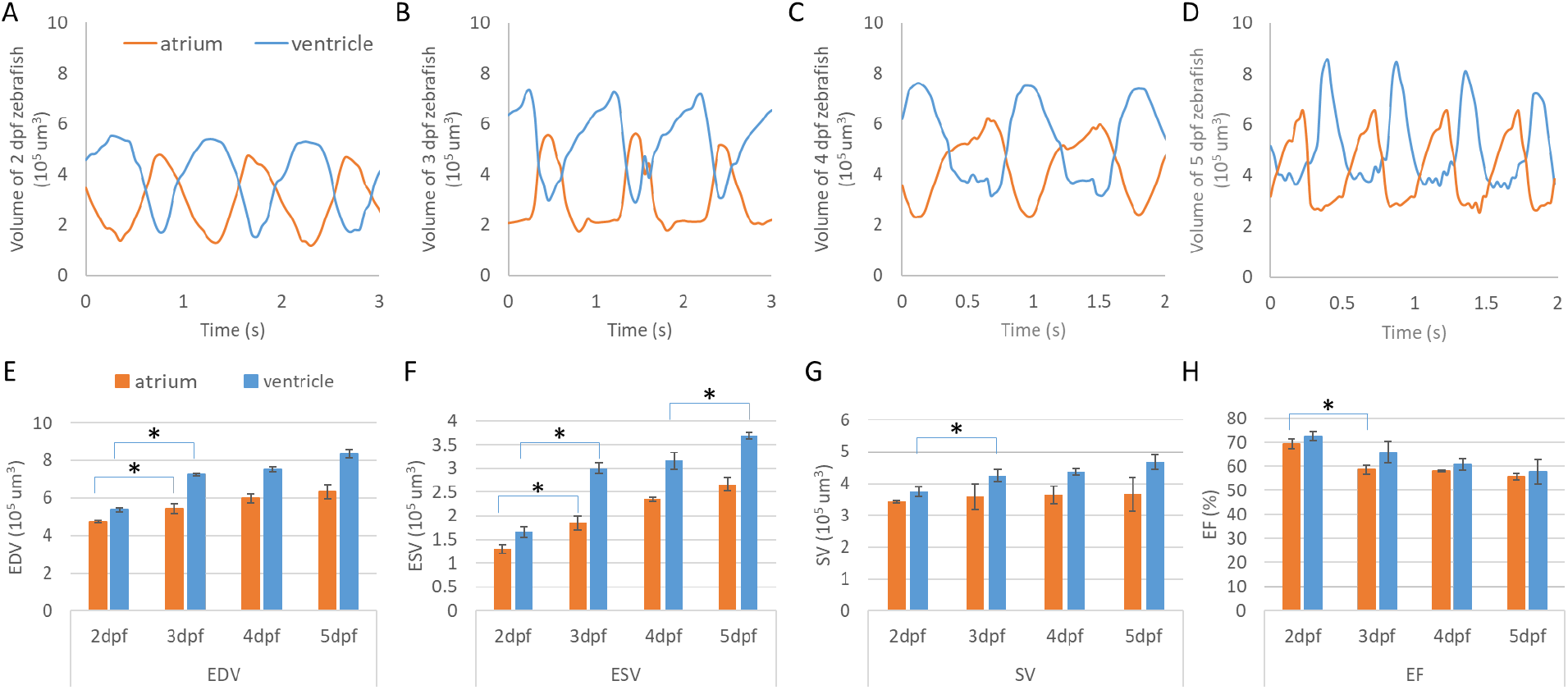
Cardiac mechanics analysis of developing zebrafish heart. (A–D) Volume change of developing zebrafish heart was measured by U-net based auto-segmentation, showing consistent increase of volume in both atrium and ventricle. (E-F) After notable morphology change after cardiac looping, EDV and ESV were increased significantly in both atrium and ventricle. At 5 dpf, ESV was also significantly increased compared to 4 dpf. (G) While SV of ventricle showed an increasing trend, atrium remained at a similar SV level from 2 dpf to 5 dpf. (H) EF analysis demonstrated a decreasing trend from high EF at 2 dpf. *p≤0.05

## 4. Discussion

Our U-net based segmentation methods successfully provided intracardiac 3D volume structure for studying the biomechanics of zebrafish hearts trained with manually segmented LSFM images. Previously created manual segmented volume suffers from tedious and taxing processes on the researcher; thus leading to a possible room for error as time requirements increase. Furthermore, inconsistency would be induced between individuals, yielding different results of the same volume for two people. Although Akerberg *et. al.* used SegNet based auto-segmentation nicely segmented zebrafish intracardiac volume and analyze the cardiac function, their application was visualized in an early stage of zebrafish heart before trabeculation [17]. We have here adopted U-net for segmentation to apply complex geometry of inner ventricular surface until 5 dpf zebrafish heart. This study relies on a fully convolutional neural network, which allows for quick segmentations with as little as five datasets. Within this experiment, our application revolved around pairing U-net with LSFM. The novelty of this particular study was the new relation between the LSFM and U-net structure, which allows for high-level sectioning capabilities of the LSFM to generate the 3D structure of the complex zebrafish heart with ease, followed by automatic morphological analyses.

The primary objective of this study was to determine if the conjunctional use of this modality-program pair provided feasible results. This objective was satisfied with a high degree of success, as seen from the Dice Similarity Correlation Coefficient having a value perceived as exceptional, showing similarity between our perceived ground truth and the auto-segmentation (**Figure 3E**). Although we have observed errors between manual and auto-segmentation, we assume the fluorescent intensity threshold point was vague when segmentation was performed manually (**Figure 3C-D**). Therefore, our auto-segmentation network could help to process more detailed cardiac mechanical analysis compared to manual segmentation. Most interestingly in our cardiac mechanics analysis, relatively rapid contraction and slower relaxation of atrium at 2 and 3 dpf transitions to slower contraction and rapid relaxation at 4 and 5 dpf (**Figure 6A-D**). At 2 and 3 dpf, zebrafish atrium pumps blood in peristaltic motion due to lack of valves while zebrafish atrium relies on impedance pumping mechanism at 4 and 5 dpf [25]. Therefore, we may observe the atrial volume change curve shifts left at 2 and 3 dpf to shifts right at 4 and 5 dpf. In addition, after cardiac looping, cardiac function significantly changed as the ventricle developed faster in size. In the tubular shape of the heart before cardiac looping, the atrium is bigger than the ventricle [19]. At 5 dpf, when active trabeculation increases contractility of ventricle, contraction of ventricle increased significantly in corroborating with previous finding (**Figure 6F**)[16]. In the interim of increasing SV of ventricle, atrial SV stayed consistent during development (**Figure 6G**).

Despite rapid process and consistent results, the experiment conducted has some mild limitations when considering the overall capabilities being used. The most important component for this study including any deep-learning-based segmentation process requires a high-quality input dataset as the output test labels are only as good as those they are trained from. In our case, using the fluorescent label zebrafish, *tg(cmlc2:gfp),* yielded overall strong results from the ventricle, but on occasion could have issues with the atria due to lower cardiomyocyte density and its location deeper in the chest, leading to more imaging issues. Circumventing this issue was not a trivial action, as exploring extensive pre-processing methods was required to be able to find the boundary of the inner volume of the heart [26]. Other limitations which could be found in the experimentation process could be issues within the imaging process, in which user and systematic errors could hinder image fidelity.

Our methods will be compatible with other fluorescent optical imaging, such as z-scanned confocal microscopy images, providing quality 3D reconstruction. Our powerful U-net based segmentation could be be a new tool for studying a variety of future biomedical research applications. To begin, the parameters of the U-net could be tweaked to increase the accuracy and precision of the program for the zebrafish hearts, this would lead to a better quality auto-segmentation, which would lead to progress in the field of mechanobiology by using computational fluid dynamics to understand the shear stress or pressure which could affect cardiac morphogenesis [2; 15].

## 5. Conclusion

Our U-net based segmentation provides a delicate method to segment the intricate inner volume of zebrafish heart during development; thus providing an accurate and convenient algorithm to accelerate cardiac research by bypassing the labor-intensive task as well as improving the consistency in the results.

## 6. Acknowledgements

This study was supported by grants from AHA 18CDA34110150 (J.L.) and NSF 1936519 (J.L.).

